# Design and Characterization of Lipid Nanocarriers for Oral Delivery of Immunotherapeutic Peptides

**DOI:** 10.1101/2022.01.27.478027

**Authors:** Xiomara Calderón-Colón, Yichuan Zhang, Olivia Tiburzi, Jialu Wang, Giorgio Raimondi, Julia Patrone

## Abstract

The use of therapeutic proteins and peptides is of great interest for the treatment of many diseases, and advances in nanotechnology offer a path toward their stable delivery via preferred routes of administration. In this study, we sought to design and formulate a nanostructured lipid carrier (NLC) containing a nominal antigen (insulin peptide) for oral delivery. We utilized the design of experiments (DOE) statistical method to determine the dependencies of formulation variables on physicochemical particle characteristics including particle size, polydispersity (PDI), melting point, and latent heat of melting. The particles were determined to be non-toxic *in vitro,* readily taken up by primary immune cells, and found to accumulate in regional lymph nodes following oral administration. We believe that this platform technology could be broadly useful for the treatment of autoimmune diseases by supporting the development of oral delivery-based antigen specific immunotherapies.

**Highlights:** **3-5 bullets, 85 char or less**

- A Design of Experiments method led the formulation of biocompatible nanoparticles
- NLC accumulate into gut-draining lymphatic tissues following oral administration
- NLC protect their antigen cargo and promote its presentation
- NLC formulation is well-suited for oral delivery of immunomodulatory agents

**Graphical Abstract:** The development of nanostructured lipid carriers containing a nominal antigen (insulin peptide) for oral delivery consists on (1) nanoparticle formulation using a statistical method, (2) *in-vitro* studies to assess cellular toxicity and uptake and T cell activation, and (3) *in-vivo* studies to assess bio-distribution.

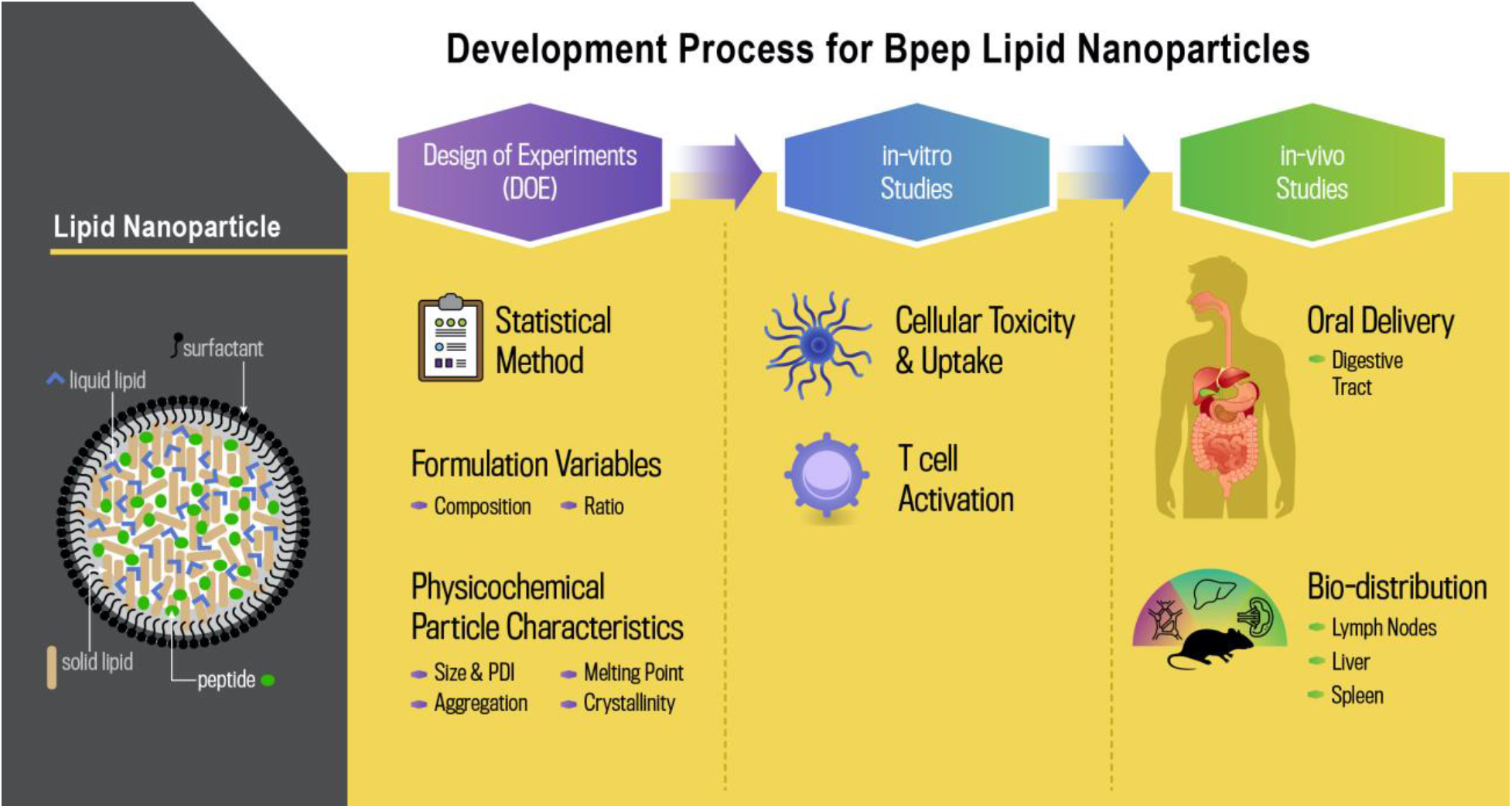

## Background

Nanoparticle-mediated delivery of therapeutically active biomolecules holds great potential for the treatment of many diseases (1-4). Over the past two decades there has been a resurgence of interest in therapeutic proteins and peptides, which can have specific advantages over small molecules due to their potential for immunomodulation, either via antigenic or adjuvant properties (5, 6). Therapeutic peptides are most often administered parenterally and require frequent doses due to their short half-life in physiological fluids (7, 8). While orally administered formulations are desirable and likely to improve patient compliance, achieving sufficient drug stability and absorption via this route remains a significant challenge. Nanoparticle technologies present an opportunity to mitigate these challenges by both increasing stabilization of therapeutic peptides and offering an approach that can target regions of the gastrointestinal tract (9). Lipid-based particles specifically, including solid lipid nanoparticles (SLN) and nanostructured lipid carriers (NLC), have shown important advantages for oral delivery in comparison to polymer-derived formulations (e.g. p*oly(lactic-co-glycolic acid) or* PLGA), due to their enhanced biocompatibility and potential for promoting intestinal absorption of active molecules (8, 10, 11). They can be made using lipids and surfactants generally regarded as safe (GRAS) by the FDA, and have been shown to be appropriate for controlled release of a wide range of active ingredients. Drug-loaded lipid nanoparticles (LNp) tend to increase the oral bioavailability of encapsulated therapeutics, due in part to a propensity for transcellular permeation of epithelial barriers, and accumulation in the lymphatic system (8, 12-16). The clinical success of therapeutic nanoformulations ultimately depends upon many factors including drug characteristics, particle physicochemical properties, and chemical composition (17). Their limited clinical translation is due in part to a lack of mechanistic understanding related to the impact of formulation variables and synthesis methods on bioavailability, targeting, and release of the drug cargo (18).

One important driver for the development of orally delivered peptide therapeutic strategies is the promising results of antigen-specific immunotherapy (ASI): the delivery of disease-associated antigens to antigen presenting cells (APC) in a tolerogenic manner to prevent or stop the immune reactivity underlying diseases (19-21). Multiple forms of ASI have demonstrated efficacy in preclinical models of prevalent autoimmune diseases like Type 1 Diabetes, Multiple Sclerosis, and Inflammatory Bowel Disease (1, 22, 23). Autoimmune diseases are caused by chronic immune responses targeting host cells and tissues, and current accepted treatments are limited to broadly immunosuppressive or anti-inflammatory strategies. Because of its specificity of immunomodulation, ASI could potentially eliminate the side effects of systemic immunosuppression. LNp platforms are a suitable choice for realizing ASI as an oral therapy that avoids first-pass hepatic metabolism by the liver, limits systemic toxicity, and localizes therapeutics into lymphoid tissues of the gastrointestinal tract, the likely physiological sites of activation of autoimmune responses (2, 5, 8).

To achieve the design of a nanoparticle formulation suitable for stable oral administration of peptides for ASI, we focused this study on the development of NLC formulated to encapsulate a mimotope of the insulin B-chain peptide sequence 9-23 (Bpep) shown to have potential for prevention of diabetes development when used in subimmunogenic conditions (24-28). Our team has previously optimized a facile and scalable process for synthesis of ultra-small SLN based on the phase inversion temperature (PIT) method (29). Use of the PIT method increases clinical translatability by allowing precise control over particle characteristics, such as chemical composition, size, polydispersity, thermal behavior, crystallinity, and synthesis scalability. We employed Design of Experiments (DOE) methodology to formulate NLC and systematically vary synthesis parameters to determine dependencies on nanoparticle physicochemical characteristics. Promising NLC formulations were evaluated for their encapsulation of Bpep, their biocompatibility, interactions with antigen-presenting cells (APC), and physiological trafficking following oral administration. Our studies confirmed the encapsulation of Bpep into biocompatible NLC, and demonstrated their rapid uptake by APCs, as well as trafficking to the regional lymph nodes. Further, the Bpep-NLCs delivered an intact peptide that enabled proper engagement of diabetogenic T cells by APC, indicating the potential of this strategy for orally-delivered ASI approaches.

## Methods

### Mice

Non-obese diabetic (NOD) mice (ShiLtJ Stock No: 001976) and TCR BDC12-4.1 (Stock No: 009377) were purchased from Jackson Laboratories and bred at the Johns Hopkins School of Medicine facility. All animal experiments were conducted in accordance with the National Institutes of Health guide for use and care of laboratory animals, and under a protocol approved by the Johns Hopkins University (JHU) Animal Care and Use Committee.

### Hydrophobic Ion Pairing

In this study, 10 μL of the insulin Bpep (100 μg total) was mixed with 90 μL of 1.5 mg/mL sodium deoxycholate (135 μg total) and incubated at room temperature for 30 minutes to allow for complex formation. The described amounts were chosen to recreate a sodium deoxycholate to insulin molar ratio of nearly 6:1, which has been previously described as the optimal amount for this pairing process (30). After complex formation, the mixture was centrifuged for 15 minutes at 14,000 rpm. Following centrifugation, the supernatant was removed and saved for further analysis, while the complexed insulin sodium deoxycholate pellet was used as the starting material for NLC formation. Despite the knowledge of the optimal ratio, in preliminary testing additional ratios of insulin to sodium deoxycholate were tested to ensure maximum pairing efficiency was achieved.

### Determination of Hydrophobic Ion Pairing Efficiency

To determine the efficiency of the hydrophobic ion pairing (HIP) process, the supernatant obtained from the protocol described above was analyzed by a bicinchoninic acid (BCA) assay to determine the amount of protein remaining in the supernatant after pairing. A Pierce BCA Protein Assay kit from ThermoFisher was used in accordance manufacturer’s recommendations. All points of the standard curve and unknown supernatant samples were tested in triplicate. Additionally, a peptide sample of known concentration was tested to ensure the BCA results were accurate.

### Processing of Nanostructured Lipid Carriers

NLCs were prepared using the PIT method, which keeps the composition constant while the temperature is changed (31, 32). This versatile method allows the synthesis of small sizes and low polydispersity lipid nanoparticles with controlled latent heat of melting and melting point (33). In this study, different experimental factors in combination with the PIT method were systematically investigated to study the effect on NLCs particle formation and physical properties of the LNp.

### Design of Experiments

In this study, the DOE processing variables were surfactant composition, solid to liquid lipid ratio, and lipid to surfactant ratio. This study included three factors (surfactant composition, solid to liquid ratio, and lipid to surfactant ratio), with levels of (BrijO10 and Gelucire 44/14), (50:50, 70:30, and 90:10), and (1:1, 1:1.5, 1:2), respectively. The responses measured were particle size, polydispersity (PDI), melting point, and latent heat of melting. The design called for 11 runs, which are listed in Table 2. The solid lipid (tetracosane (C24), liquid lipid (tocopherol), processing temperature (70 °C), and peptide concentration were held constant. In this work, the DOE-Custom Designer platform from the commercial statistical software JMP (SAS Institute, Cary, North Carolina) was used to generate a Resolution V Design.

**Table 1.** Experimental conditions for three different pairing ratios prior to particle synthesis.

| Condition | Amount Peptide (μg) | Amount Sodium Deoxycholate (μg) | Molar Ratio(sodium deoxycholate:insulin) | Calculated Pairing Efficiency |
| --- | --- | --- | --- | --- |
| 1 | 100 | 45 | 1.8 | 82% |
| 2 | 100 | 135 | 5.4 | 93% |
| 3 | 100 | 270 | 10.7 | 91% |

**Table 2.**
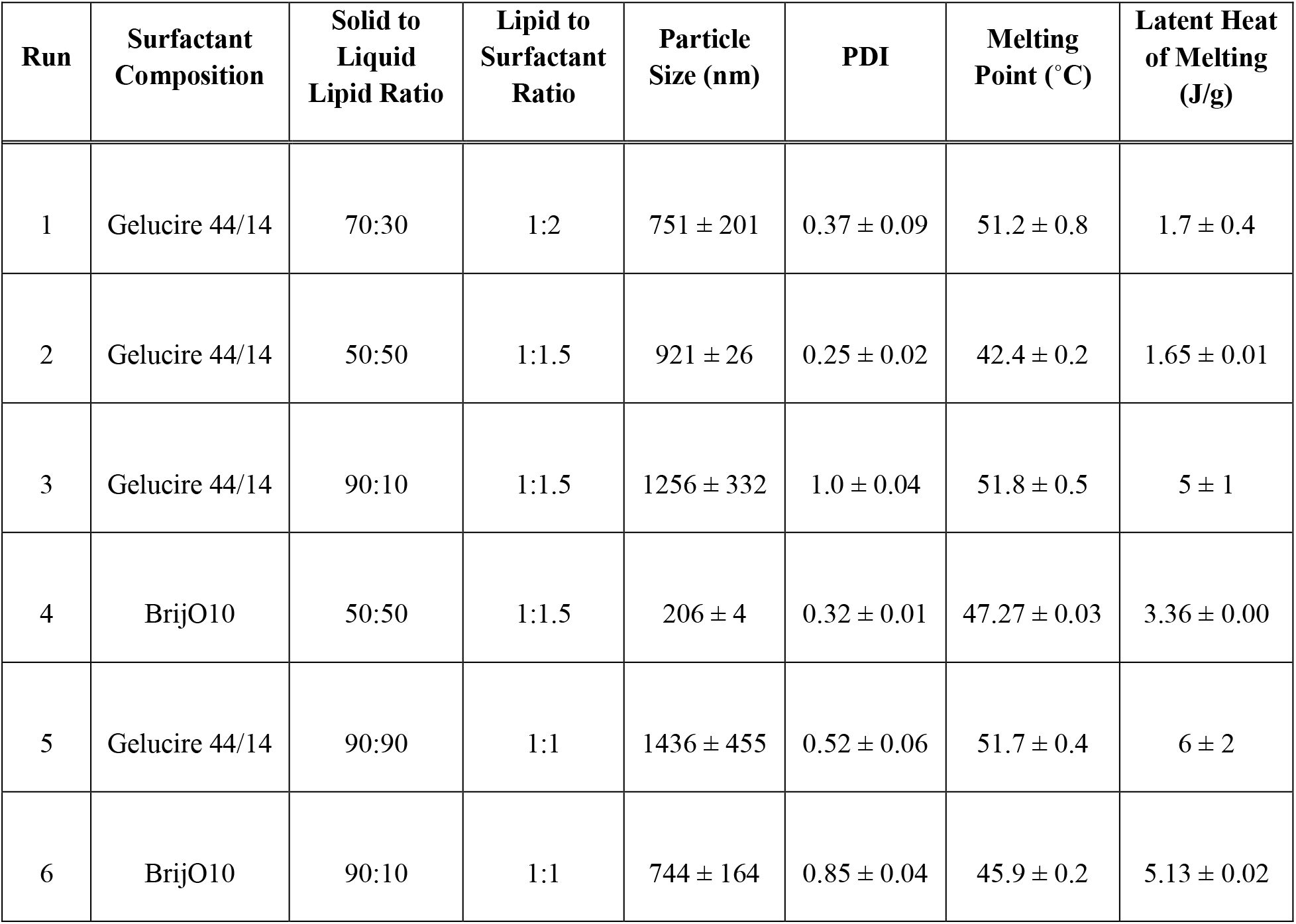
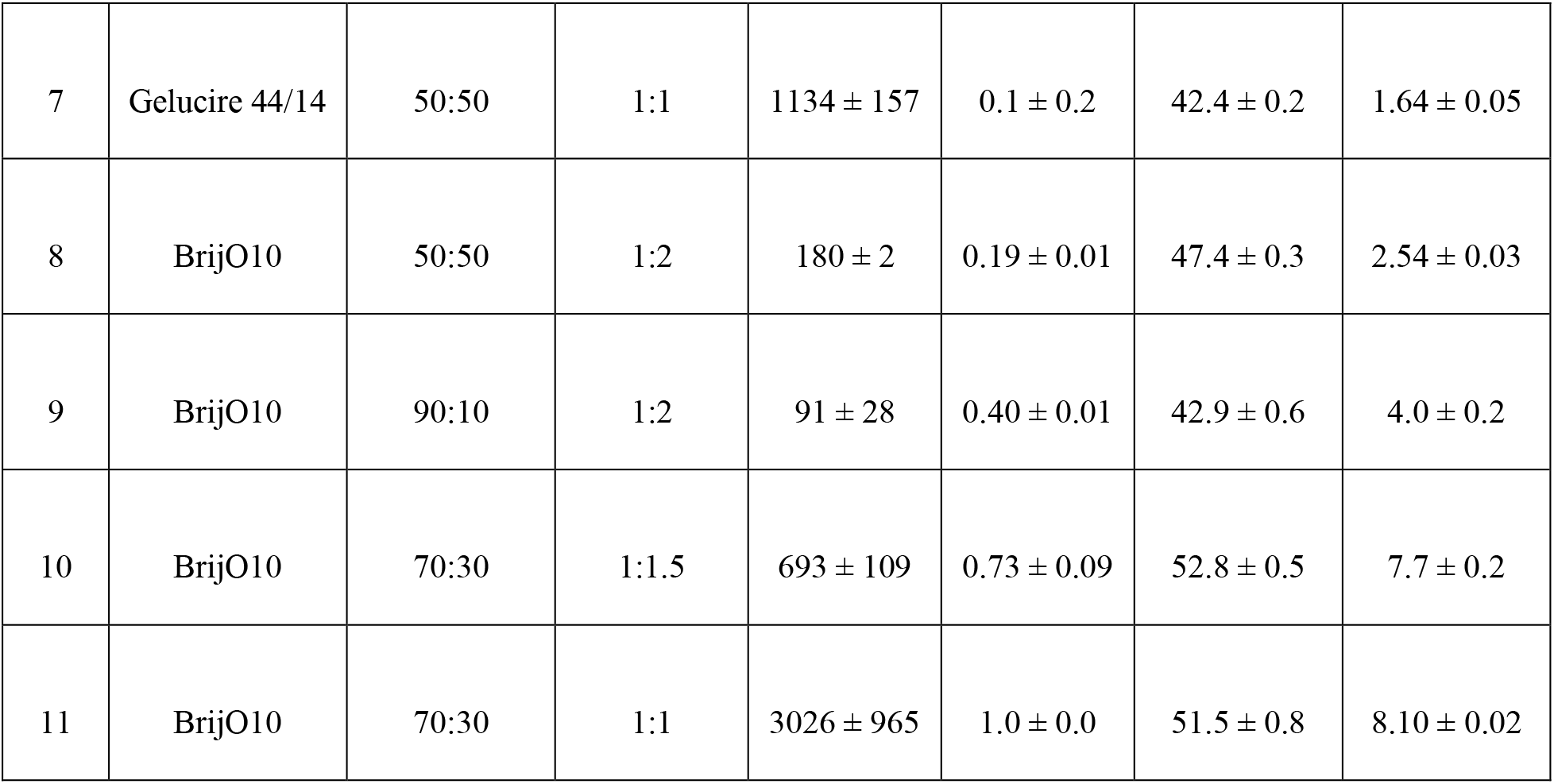
Summary of the processing parameters for the formation of Bpep-NLCs and DOE results including values for each of the responses (particle size, PDI, melting point, and latent heat of melting).

**Table 3.** Summary of DOE statistical analysis including range achieved for each response, influential factors, and confidence level.

| Response | Range Achieved | Influential Factors | Confidence Level (%) |
| --- | --- | --- | --- |
| Particle Size | 91 – 3000 nm | Lipid to Surfactant Ratio | 99.99 |
|  |  | Solid to Liquid Lipid Ratio | 99.98 |
| PDI | 0.1 – 1.0 | Surfactant Composition | 99.99 |
|  |  | Lipid to Surfactant Ratio | 99.99 |
|  |  | Solid to Liquid Lipid Ratio | 99.97 |
| Melting Point | 42.4 – 52.8 °C | Lipid to Surfactant Ratio | 99.99 |
|  |  | Solid to Liquid Lipid Ratio | 99.99 |
| Latent Heat of Melting | 1.31 – 8.10 J/g | Surfactant Composition | 99.99 |
|  |  | Lipid to Surfactant Ratio | 99.99 |
|  |  | Solid to Liquid Lipid Ratio | 99.99 |

### HLDI-loaded NLCs

HLDI-loaded NLCs were prepared starting with the fluorescent dye (0.35 mg/mL final concentration) and ethanol (60 μL) combined into a vial. Following solubilization, the surfactant (0.80 g of Gelucire 44/14) was added into the vial, co-melted at 90 °C and stirred. Subsequently, the lipids (0.20 g of Tetracosane (solid lipid) and 0.20 g of Tocopherol (liquid lipid)) were added.

The mixture was again co-melted at 90 °C and stirred. Lastly, DI water was added to the mixture, and heated. The resulting mixture was stirred using a vortexer while allowing it to cool, promoting the creation of a nanoemulsion. In parallel, a nanoemulsion without fluorescent dye was prepared (to be used as a control). In addition, in some experiments a HLDI solution was implemented as control sample. The HLDI solution was prepared by adding HLDI fluorescent dye in ethanol to obtain a dye concentration of 0.35 mg/mL.

### Particle Size and Polydispersity

The particle size and PDI of the NLCs were measured using a Malvern Zetasizer Nano S dynamic light scattering system. In this technique, the frequency of the shifted light is used to measure the particle size. The particle size of the NLCs was measured after adding 25 μL of particles into 3 mL of water. The water was filtered with a 0.2 μm filter directly into a cuvette prior the addition of the NLCs. The particle diameter and PDI of particle size distribution are reported. Each formulation of the DOE was measured three times.

### Melting Point and Latent Heat of Melting

The thermal behavior of NLCs was studied using a Mettler-Toledo differential scanning calorimeter (DSC) equipped with an auto-sampler and liquid nitrogen as the cooling source. The as-prepared NLCs were pipetted into a 40 μL aluminum pan with a mass of approximately 25 mg. The aluminum pan was then hermetically sealed to minimize moisture loss during the DSC scan. The NLCs were measured from 5 to 80 °C. The latent heat of melting was calculated using the integral of the area under the curve (amount of energy) divided by the amount of material. The valley in the DSC plot represents the melting point. The melting point and the latent heat of melting are reported.

### Particle Size Optimization and Stability

To determine the optimal NLC composition with regards to both particle size and stability, a small scale stability study was conducted. Three replicates of the optimized peptide NLC formulation were prepared, and the particle size of each formulation was measured using the Malvern Zetasizer Nano S dynamic light scattering system right after generation and then weekly for a total for four weeks.

### Transmission Electron Microscope

A small amount of the NLC suspension was pipetted onto a special cryo-TEM grid that was secured to the cryo-plunger (Gatan Cryoplunge TM3). The grid was plunged into liquid nitrogen to produce a thin amorphous ice (glass-like solid) that preserves the microstructure in hydrated state. The plunged TEM grid was immediately transferred onto a cryo-TEM holder (Gatan 914) using Gatan Cryotransfer System, and then loaded into the TEM. The entire process during specimen preparation through TEM observation was kept in liquid nitrogen temperature (lower than -173 °C). TEM investigation of cryo-samples was performed by using a JEOL transmission electron microscope (JEM 2100 LAB6 TEM) at 100 kV.

### Encapsulation Efficiency

The encapsulation efficiency of the peptide into the NLC formulation was determined via the CBQCA protein quantitation kit (Thermo Fisher Scientific). Three separate NLC peptide formulations were created using the optimized formula previously determined in the DOE. Each of the NLC preparations were separated into an aqueous and organic phase by placing a known volume in a 100 MWCO spin column and centrifuging for 1.5 hours at 3500xg. Post centrifugation, the filtrate is the aqueous phase and the retentate is the organic phase. The total volume of each phase was noted. Samples (100 μL in triplicate) from the aqueous, organic and unprocessed NLC formulation were input into the CBQCA assay protocol to determine the amount of peptide present. A standard curve was created using a stock of the peptide to help ensure accurate interpolation of the unknown samples. Upon completion of the CBQCA assay, the interpolated concentration of each phase (aqueous and organic) and the total volume of each phase were used to back calculate the total amount of peptide in each portion of the overall NLC formulation. Peptide found in the organic phase was considered to be encapsulated into the NLC, while the peptide in the aqueous phase was considered as free, un-encapsulated peptide. Therefore encapsulation efficiency was determined as the amount of peptide in the organic phase divided by the total amount of peptide, defined as the sum of the peptide found in both the aqueous and organic phase. The three replicate formulations were created and tested on three separate days.

### Cellular Toxicity and Particle Uptake

Mouse bone marrow-derived dendritic cells (BMDC) were generated from NOD mice according to a previously published protocol (34). For assessment of cellular toxicity, differentiated BMDCs were cocultured overnight with unloaded NLC (formulations indicated in the text) at different concentrations (0.625 μg/mL, 1.25 μg/mL, 2.5 μg/mL, 5 μg/mL, 10 μg/mL, and 20 μg/mL), and then stained with Fixable Viability Dye eFluor 80 (eBioscience/ThermoFisher) and anti-CD11c antibody (clone N418, eBioscience). For the assessment of particle uptake, BMDCs were cocultured with 1 μg/mL HLDI-loaded NLC at 1 μg/mL for 10 to 30 minutes at 37°C, and then stained with anti-CD11c. Cell fluorescence was measured on a BD LSRII flow cytometer (BD Bioscience) and analyzed using FlowJo software (version 10.5.3, TreeStar).

### Biodistribution

HLDI-NLC (15 mg particle mass/animal) suspended in 200 μL of sterile saline was administrated to NOD animal as a single oral gavage. Lymphoid tissues including spleen, inguinal lymph nodes (iLNs), mesenteric lymph nodes (mLNs), and pancreatic lymph nodes (pLNs) were extracted from treated and untreated (control) animals four hours after administration. The fluorescence signal accumulated in these tissues was detected via *in vivo* imaging system (IVIS) spectrum imaging system (Perkin Elmer), with excitation set at 675 nm and emission at 760 nm.

### T Cell Isolation

CD4+ T were isolated as previously described (35). Briefly, Spleen and lymph nodes were extracted from mice and processed to generate single cell suspensions. CD4+ T cells were then enriched via negative selection using Dynabeads Untouched Mouse CD4 T cells Kit (Thermo Fisher Scientific) according to manufacturer protocol.

### T Cell Proliferation Assay

On day seven of culture, BMDCs were cocultured with either free mimotope (0.04 μg/mL) or Bpep-NLC (100 μg/mL) and then stimulated overnight with LPS (200 ng/mL).The following day, cells were collected and cocultured for four days with T cells that had been isolated from BDC12-4.1 transgenic mice and stained with CFSE as previously described (34). T cell proliferation (dilution of CFSE fluorescence intensity) was measured via flow cytometry and data analyzed via ModFit LT™ software.

### ELISpot Assay

BMDCs were incubated, during the last day of differentiation, with either free Bpep (1 μg/mL), Bpep-NLC (125 μg/mL), or control NLC (125 μg/mL) and then stimulated overnight with Poly(I:C) (20 μg/mL, Sigma Aldrich). T cells from BDC12-4.1 mice were enriched as described above and cocultured with the different BMDCs groups in anti-mouse IFN-γ (R4-6A2, eBioscience) coated 96-well filtration plate (Millipore Sigma) for two days. Secondary biotinylated antibody IFN-γ (RA-6A2, eBioscience) was added on the third day, and streptavidin-horseradish peroxidase the day after. The plate was then read using a Zeiss KS EliSpot reader.

### Statistical Analysis

All values are reported as mean ± SD. Differences in ELISpot were assessed using two-tailed unpaired Student’s t-test. All analysis was performed using Graph Prism 7.0a version (GraphPad, San Diego, CA). A p-value <0.05 was considered statistically significant.

## Results

### NLC Formulation, Toxicity, and Cellular Uptake

The overall goal of this study was to define NLC formulation parameters to achieve maximal, stable loading of Bpep for effective oral delivery. Based on preliminary efforts by our laboratories to design and formulate NLC for the encapsulation of a small molecule (tofactitinib, a Jak inhibitor), we first performed a screening DOE to determine which NLC formulation variables resulted in a nanoemulsion using specific combinations of liquid and solid lipids as well as two surfactants (data not shown). Following down-selection of formulations that resulted in successful nanoemulsions, we then evaluated whether the surfactant type or a variation in lipid to surfactant ratio would present different degrees of cellular toxicity. We tested three different formulations (Brij 1:1, Gelucire 1:2, and Brij 1:2) by measuring cell viability after overnight co-culture of mouse bone marrow derived dendritic cells (BMDC – an *in vitro* representation of the antigen presenting cells that would be targeted *in vivo*) with titrations of NLCs. Formulation Brij 1:2 (containing BrijO10 surfactant and a 1:2 lipid to surfactant ratio) was found to be fairly toxic, with appreciable loss of cell viability at a concentration greater than or equal to 1.25 μg/mL (Figure 1). Formulation Brij 1:1 (also containing BrijO10 surfactant but at a 1:1 lipid to surfactant ratio) presented noticeable toxicity starting at a concentration of 2.5 μg/mL. Formulation Gelucire 1:2 (containing Gelucire surfactant at a 1:2 lipid to surfactant ratio) did not present any significant negative impact on DC viability at any of the concentrations tested. We repeated this assessment for NLC Formulation Gelucire 1:2 and found no appreciable toxicity at concentrations as high as 500 μg/mL (data not shown). Based on these results, we then formulated the NLC to encapsulate the far red dye 1,1’,3,3,3’,3’-Hexamethylindodicarbocyanine Iodide (HLDI-NLC) as a means to assess the degree of cellular uptake. BMDC were co-cultured with HLDI-NLC at 1 μg/mL for 10 or 30 minutes, washed, and the intensity of HLDI fluorescence incorporated by CD11c+ BMDC determined via flow cytometry. The results indicated a rapid and progressive uptake of the Gelucire 1:2 NLC by BMDC (Figure 1C).

**Figure 1.**
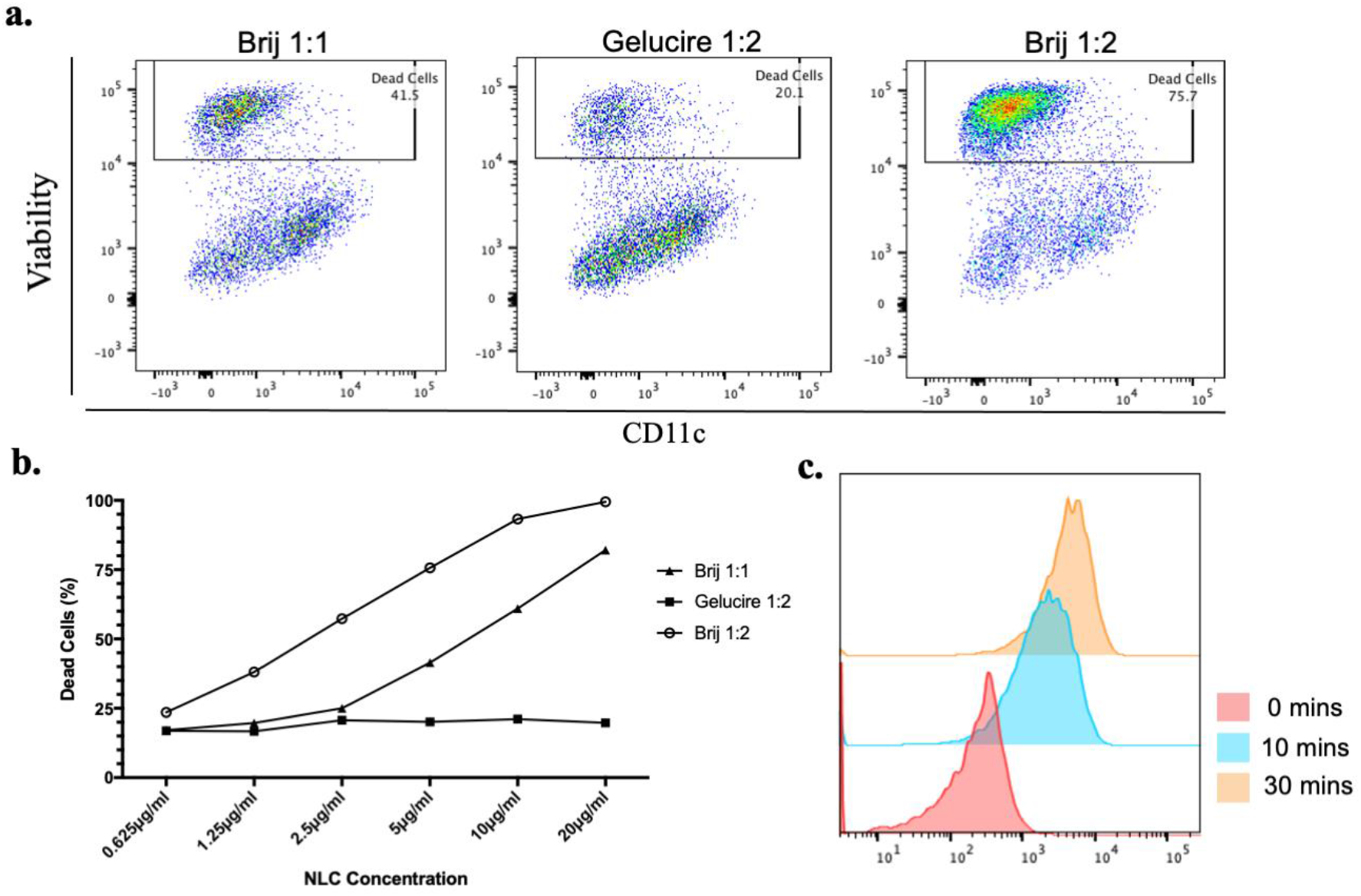
Cellular toxicity and uptake of NLC formulations. **a.** BMDC were cocultured overnight with different NLC formulations at multiple concentration. The toxicity of NLCs was determined by measuring the viability of BMDCs via flow cytometry. Representative results with formulation Brij 1:1, Gelucire 1:2, and Brij 1:2 at 5 μg/ml, with death rate 41.5%, 20.1%, and 76.7% respectively. **b.** Correlation between cell death and concentration of different formulation of NLC in BMDC as in **a**. **c.** Cellular uptake; Gelucire 1:2 NLC were synthesized containing the dye HLDI and cocultured with BMDCs for the indicate time. Uptake was determined by co-staining of HLDI and CD11c via flow cytometry.

### Hydrophobic Ion Pairing Efficiency

Following the screening DOE and down-selection of NLC formulations with low toxicity, we then sought to encapsulate Bpep using the PIT method, as before. However, despite our preliminary successful encapsulation of tofacitinib, our efforts to encapsulate the peptide resulted in a low encapsulation efficiency (data not shown). Therefore it was necessary to increase the liposolubility of the peptide prior to incorporation into the nanoformulation. We explored HIP, which involves complexing the target molecule with a secondary molecule (typically of the opposite charge). This interaction masks the initial charge of the target molecule and subsequently alters the solubility properties (36). This technique has been used previously to increase the liposolubility of insulin and works by complexing the insulin peptide with sodium deoxycholate (30). Initial testing of the HIP efficiency indicated that 82 – 93% of the 100 μg of peptide was successfully paired (**Error! Reference source not found.**). Condition 2 was determined to be optimal and was selected for all subsequent experiments. A BCA assay was used to confirm that the pairing efficiency was approximately 93% prior to LNp synthesis.

### Design of Experiments

To further optimize NLC properties for incorporation of the peptide cargo and gain a deeper understanding of the influence of surfactant/lipid type and lipid to surfactant ratio on particle physicochemical properties, we next utilized a higher resolution DOE. Table 2 lists the processing parameters included in this DOE statistical analysis: surfactant composition, solid to liquid lipid ratio, and lipid to surfactant ratio. For each run we measured four responses: particle size, PDI, melting point, and latent heat of melting. A summary of these responses, the most influential factors, and corresponding confidence level from the statistical analysis are shown in **Error! Reference source not found.**. To maximize cell interaction (uptake and trafficking), we sought to achieve a small particle size with low polydispersity, and low toxicity. The synthesized Bpep-NLCs ranged in size from 91 to 3000 nm, where the smallest NLCs were synthesized using BrijO10 as a surfactant, using a solid to liquid lipid ratio of 90:10, and a lipid to surfactant ratio of 1:2.

The interaction of the lipid to surfactant ratio and solid to liquid lipid ratio were found to be statistically significant factors for all the responses tested. In addition, surfactant composition was significant in impacting the PDI and latent heat of melting. The results of the DOE further demonstrated that depending on the processing conditions, the resulting Bpep-NLCs had a melting point between 42.4 and 52.8 °C and a latent heat of melting from 1.31 to 8.10 J/g, confirming that the nanoparticles will maintain their characteristics at body temperature. The solid to liquid lipid ratio and the lipid to surfactant ratio are influential for determining the melting point of the Bpep-NLCs.

### Particle Size Optimization and Stability

Following the toxicity testing and higher resolution DOE, we utilized the statistical analysis outputs to further guide Bpep-NLC optimization to achieve particles <100 nm in diameter. Based on the DOE response showing that particle size is influenced by the lipid to surfactant ratio (Figure 2A, P < 0.0001), we then varied the solid to liquid lipid ratio incrementally while keeping all other experimental factors constant to measure the impact on particle size and PDI. As displayed in **Error! Reference source not found.**B, Bpep-NLC formulations with a lower solid lipid percentage yielded smaller particles that were very uniform, indicating the importance of the liquid lipid on particle size. Three replicates of the 50:50 solid lipid to liquid lipid ratio Bpep-NLC formulation were then prepared side-by-side and we monitored their particle size and stability over time (Figure 2C). The particle size remained consistent across the three replicates throughout the study, with only a small increase in week 4, indicating the stability of this formulation. To further investigate the size and shape of the lipid nanoparticles, we characterized them by cryo-TEM (**Error! Reference source not found.**3). This image shows the presence of rounded nanoparticles and validated the size results previously obtained by DLS.

**Figure 2.**
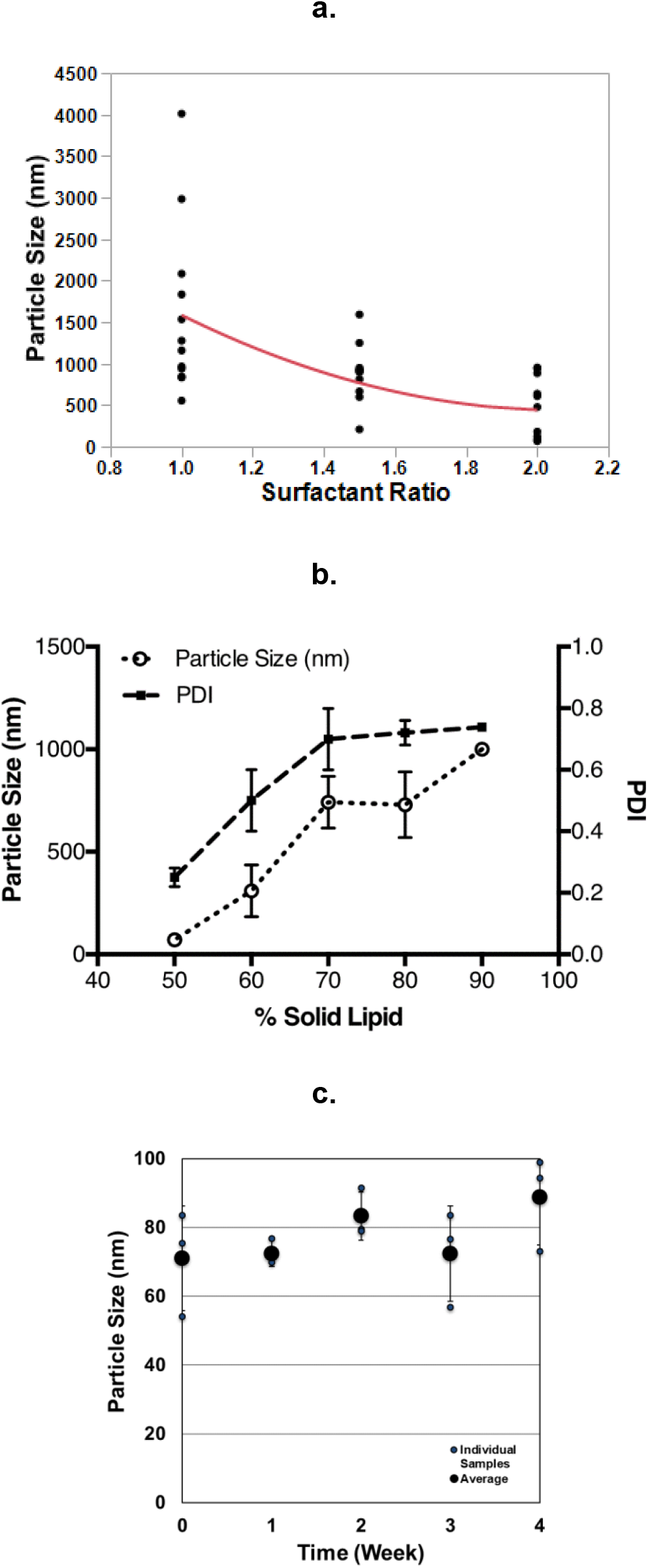
Measurement of size, PDI, and stability of Bpep-NLC. a. The relationship between surfactant ratio and particle size (P < 0.0001). b) Particle size and PDI as a function of solid lipid. c) Particle size as function of time.

**Figure 3.**
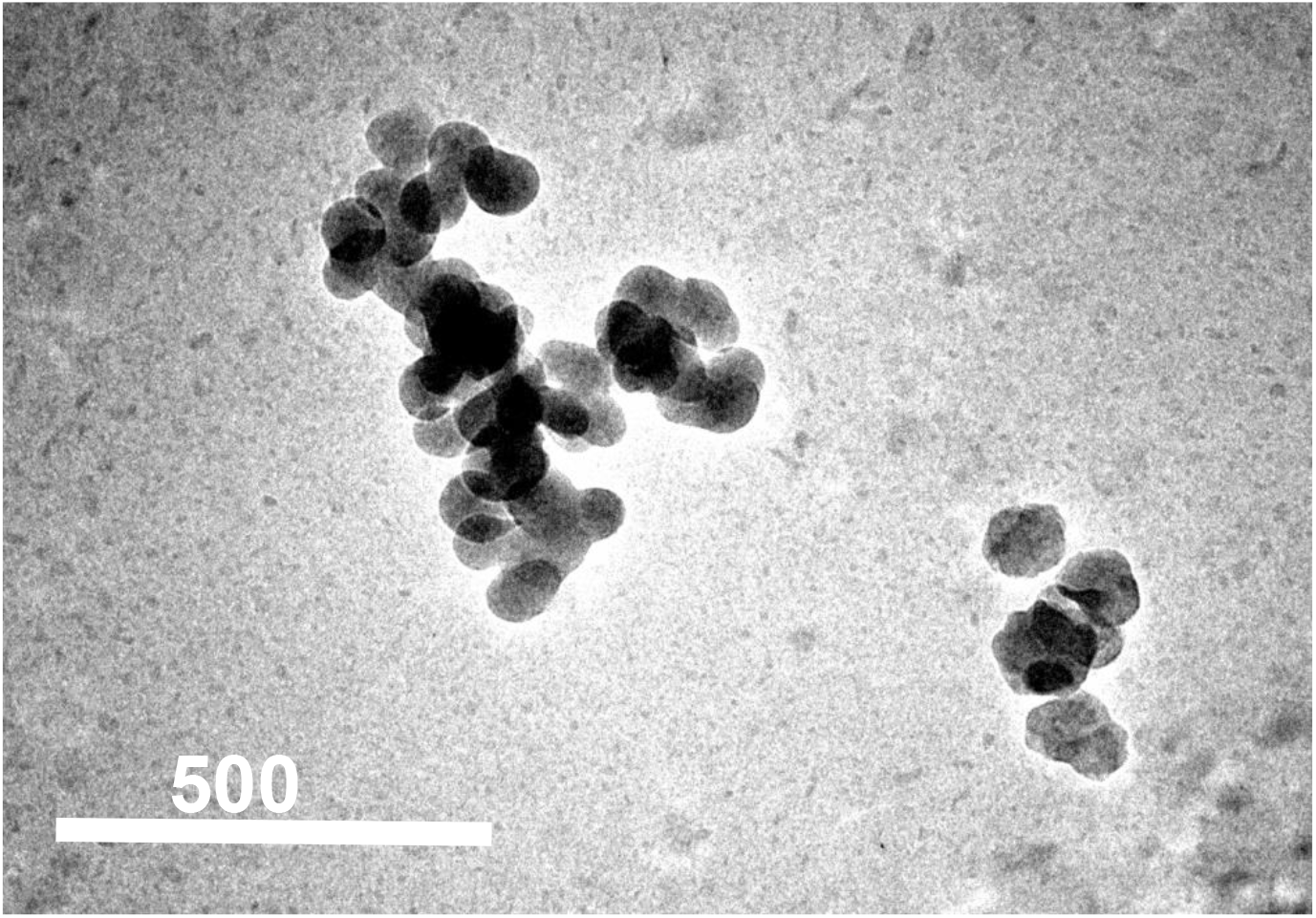
Top view TEM image of control (empty) NLC prepared using the PIT method.

### Encapsulation Efficiency

To quantify the peptide encapsulation efficiency (EE%), three identical Bpep loaded NLC formulations were created and processed via spin column centrifugation to separate out the aqueous and organic phases. To measure the amount of Bpep in each portion, we employed the CBQCA assay, a method that is not affected by the presence of lipids or detergents, making it ideal for studying NLC formulations. The results from this experiment are displayed in Table 4, revealing an average Bpep EE% of 78%.

**Table 4:** Insulin B-peptide Encapsulation Efficiency Results.

| Replicate | Encapsulation Efficiency |
| --- | --- |
| **1** | 72% |
| **2** | 83% |
| **3** | 79% |
| **Average** | 78% ± 5% |

### Biodistribution

As our goal was to engineer a delivery vehicle for the accumulation of disease-specific peptides in lymphoid tissues following oral administration, we tested if the selected formulation could cross the intestinal barrier and accumulate in draining lymphoid tissues. For this study, we administered a bolus of HLDI-NLC to NOD mice via oral gavage and, four hours later, determined the intensity of HLDI fluorescence emitted by draining and nondraining lymphoid tissues via *in vivo* imaging system (IVIS) analysis. Our results (Figure 4) indicated that, following oral administration, NLCs (formulation Gelucire 1:2) accumulated in pancreatic and mesenteric lymph nodes, and the spleen, but not in inguinal lymph nodes (non-draining tissues), as expected.

**Figure 4.**
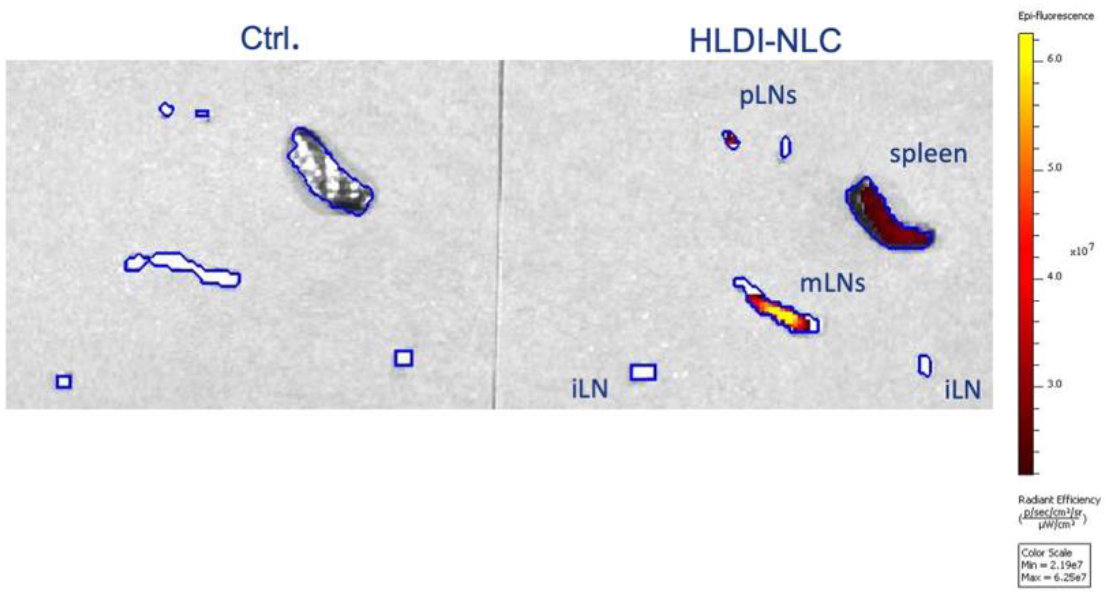
NLC Biodistribution. HLDI-NLCs were orally administrated into NOD mice. Tissues of interest (shown in graph) were extracted 4h later and the tissue-specific accumulation was detected using IVIS. Comparing with control mice, our formulation of NLCs tend to accumulate into gut-draining lymphatic tissues (mLNs, and pLN) and to a certain degree to the spleen.

### Antigen Integrity and Presentation by APC

Having confirmed the rapid uptake by APCs and the *in vivo* distribution of our selected NLC formulation, we examined if the process of encapsulation would preserve peptide integrity and if the particulate product could be processed by APCs for successful presentation to T cells. To this end, we used insulin-specific BDC12-4.1 TCR transgenic mouse CD4+ T cells, as they can be activated by APCs presenting the insulin peptide mimotope employed in this study (37, 38). We first examined the proliferation of CFSE-stained BDC12-4.1 T cell induced by BMDC that had been pre-exposed to Bpep-NLC or free Bpep (as control). The concentration of Bpep-NLC and free Bpep were chosen to provide a theoretical equivalent amount of mimotope to APCs in each condition. Figure 5A shows that BMDC preincubated with Bpep-NLC induced a degree of T cell proliferation equivalent to that of DC loaded with free peptide: a confirmation of cargo integrity and proper processing of Bpep-NLC by APCs. We also tested if the presentation of Bpep carried by NLC would induce effector activities in the recognizing T cells. To this end, we measured via ELISpot assay the number of IFN-γ producing BDC12-4.1 T cells (an indicator TH1 effector skewing) induced by the co-culture with BMDC that had been pre-exposed to Bpep-NLC (or pre-exposed to free peptide as positive control) followed by maturation with the adjuvant Poly(I:C). Bpep-NLC loaded DC led to a number of IFN-γ producing cells similar to that of free mimotope exposed DC, while control empty NLC did not provide appreciable cytokine production (Figure 5B-C). Altogether, these results indicate that our selected formulation of NLC carries an intact peptide cargo that can be processed for successful antigen presentation by APCs.

**Figure 5.**
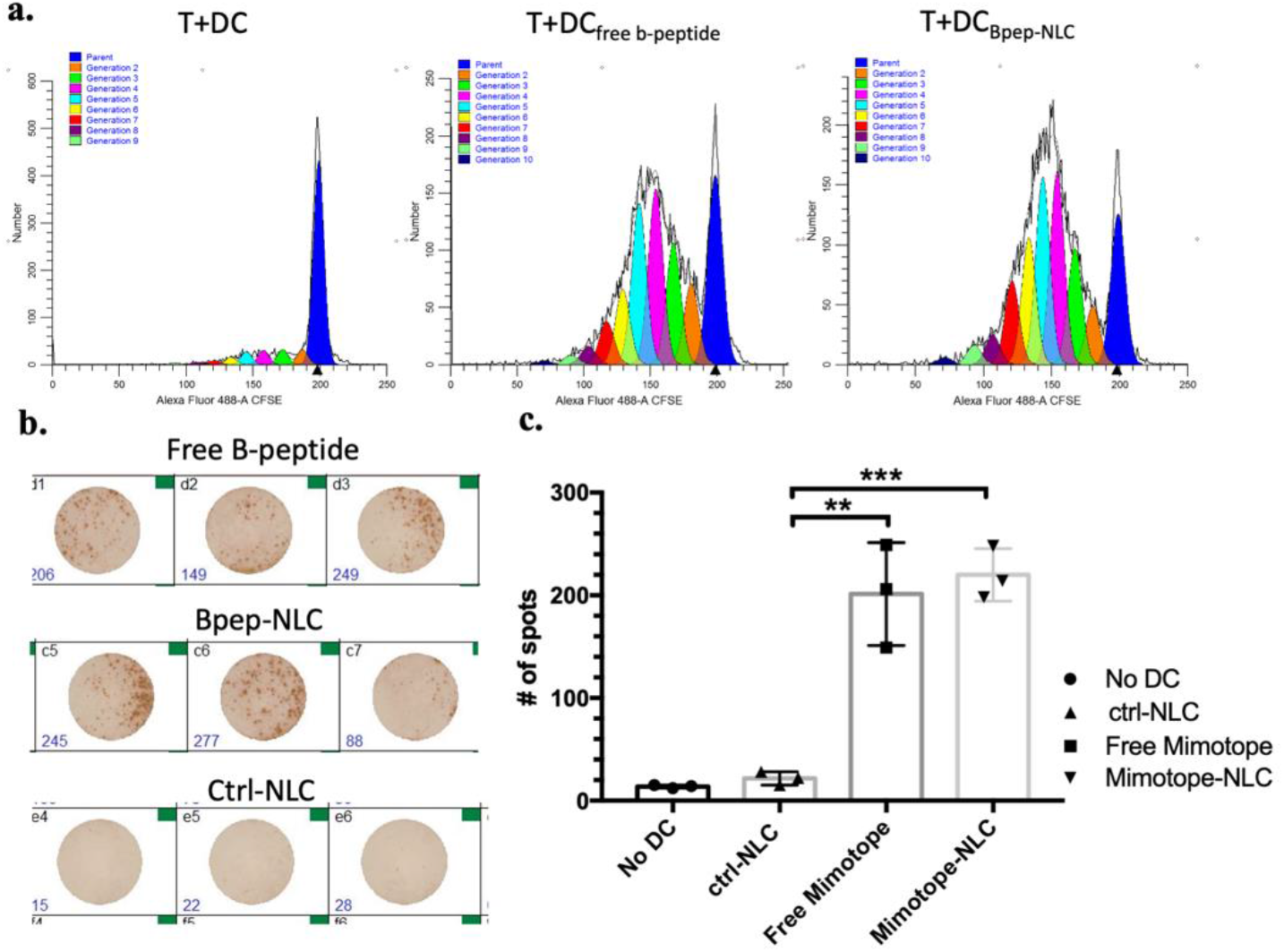
NLC encapsulation maintains antigen integrity and presentation by APC. **a**. Representative results of induction of proliferation of insulin-specific BDC12-4.1 T cells after coculture with BMDC pre-loaded with peptide of Bpep-NLC. **b**. Representative image of ELISpot assay reflecting the production of IFN- ⎕ by BDC12-4.1 T cells co-cultured BMDC pre-exposed to B-peptide or Bpep-NLC. **c**. Cumulative results from IFN-⎕ ELISpot assay described in b. n=3 independent experiments, and expressed as the mean ± SD. Statistical differences between groups are indicated as **p <0.01 and ***p <0.001

## Discussion

One of the most challenging drug delivery strategies is the oral delivery of therapeutic peptides and proteins. However, new nanoparticle synthesis techniques and formulations offer the potential to design solutions that shield the drug cargo from the harsh gastrointestinal tract environment, thus improving bioavailability, cell uptake, and permeation across epithelial barriers. Oral delivery of peptide antigens in a controlled and targeted fashion can be used toward immunotherapeutic approaches, including for the treatment of autoimmune diseases. In this study, we utilized the DOE method to design an LNp formulation to i) encapsulate a model peptide antigen and ii) target the draining lymph nodes of the gastrointestinal tract.

To reproducibly achieve a stable nanoemulsion, comprised of small, uniform particles, we systematically varied several formulation factors and performed particle analysis to confirm the impact on physicochemical properties. The benefit of using this DOE process is the ability to vary multiple parameters simultaneously and determine the statistically significant effects. Overall we found that lipid to surfactant and solid to liquid lipid ratio are key for tailoring the physicochemical properties of the Bpep loaded NLC formulations across all the responses tested. Moreover, the modulation of the solubility of the peptide allowed its incorporation into a LNp platform with a high encapsulation efficiency and stability.

DOE methods are useful to narrow particle formulation variables, however responses are somewhat limited to characteristics that can be determined using physical methods (e.g. DLS, DSC, etc.) The influence of the down-selected formulation in biological systems must therefore be determined empirically using *in vitro* and *in vivo* models. One aspect that is often neglected in the design of NLC formulations is their potential inherent toxicity – a major concern for their clinical applicability. NLCs are composed of lipid and surfactants, and there have been very few reports assessing the correlation between particle components and cellular toxicity (39-41), if we exclude the intended toxicity toward cancer cell lines. In addition to delineating the dose-response profile for key NLC candidate formulations, we identified Geluire 44/14 as a surfactant with very low toxicity, making it suitable across a wide range of applications. This characteristic facilitates flexibility in dosing, allowing variation to achieve the desired therapeutic concentration without elevated risk of systemic toxicity.

Our data indicate that our NLC formulation is well-suited for oral delivery of immunomodulatory agents, as the particles tend to accumulate into gut-draining lymphatic tissues such as mesenteric and pancreatic lymph nodes in mice. This property has been often inferred for lipid nanoparticles, but mostly investigated via laparoscopic injection rather than oral gavage (a better representation of clinical use) (42, 43). Although the mechanism(s) behind the biodistribution after oral administration is still not clear, the successful delivery of bioactive antigens that can be presented by antigen presenting cells and modulate T lymphocyte responses makes our formulation of NLC an appealing oral delivery vehicle. Not only would these features allow targeted delivery of a drug to treat gut-related immunological diseases with minimal systemic exposure, but it could also allow exploitation of the inherent tolerogenicity of gut lymphoid tissues to modulate immune reactivity to other pathologies (16, 44).

In our study, we used a derivative of an insulin peptide as proof-of-principle for the feasibility to use NLC to carry modulatory antigens to lymphatic tissues via oral administration. Our results confirmed that the peptide can be successfully encapsulated in our NLC formulation and is retained in an intact form that can be processed and presented by antigen presenting cells. Our next step will be assessing the level of *in vivo* delivery following oral administration and determining its therapeutic impact on diabetes development. The insulin mimotope was found effective when used to ‘negatively vaccinate’ mice via daily subcutaneous injections administered for 14 days (45, 46). Proving that a similar result can be achieved via oral administration would be a very important therapeutic milestone.

Every autoimmune disease is characterized by reactivity against multiple antigens which are being progressively identified. We believe that NLC represent a versatile platform technology that can be exploited to implement this growing knowledge for the actuation of antigen specific immunotherapy via administration of key antigens. Enhanced oral bioavailability and lymphoid targeting of peptides or proteins, probably combined with immune-modulatory drugs (that could be delivered via the same particles), could reasonably form the foundation of innovative and highly effective immunotherapies for many diseases.

## Abbreviations

SLNs: Solid Lipid Nanoparticles
NLCs: Nanostructured Lipid Carriers
PLGA: Poly(lactic-co-glycolic acid)
GRAS: Generally Regarded As Safe
FDA: Food and Drug Administration
T1D: Type 1 Diabetes
ASI: Antigen-specific Immunotherapy
APC: Antigen Presenting Cells
LNp: Lipid Nanoparticle
PIT: Phase Inversion Temperature
DOE: Design of Experiments
MWCO: Molecular Weight Cut-Off
HLDI: 1,1′,3,3,3′,3′-Hexamethylindotricarbocyanine iodide
HPLC: High Pressure Liquid Chromatography
BCA: Bicinchoninic Acid
HIP: Hydrophobic Ion Pairing
PDI: Polydispersity
DSC: Differential Scanning Calorimeter
CBQCA: (3-(4-carboxybenzoyl)quinoline-2-carboxaldehyde)
NOD: Non-Obese Diabetic
BMDC: Bone Marrow-Derived Dendritic Cells
iLNs: inguinal Lymph Nodes
mLNs: mesenteric Lymph Nodes
pLNs: pancreatic Lymph Nodes
EE: Encapsulation Efficiency
IVIS: *In Vivo* Imaging System

## Acknowledgements

This work was supported by grant 2-SRA-2016-310-S-B from the Juvenile Diabetes Research Foundation.

